# ML-driven design of 3’ UTRs for mRNA stability

**DOI:** 10.1101/2024.10.07.616676

**Authors:** Alyssa K. Morrow, Ashley Thornal, Elise Duboscq Flynn, Emily Hoelzli, Meimei Shan, Görkem Garipler, Rory Kirchner, Aniketh Janardhan Reddy, Sophia Tabchouri, Ankit Gupta, Jean-Baptiste Michel, Uri Laserson

**Author notes:** The author contributed equally to this paper (Alyssa K. Morrow, Ashley Thornal, Elise Duboscq Flynn, Emily Hoelzli, Meimei Shan). Formerly at Patch Biosciences (Alyssa K. Morrow, Ashley Thornal, Elise Duboscq Flynn, Emily Hoelzli, Görkem Garipler, Rory Kirchner, Aniketh Janardhan Reddy, Jean-Baptiste Michel, Uri Laserson). Currently at OverT Bio (Ashley Thornal). Currently at Neptune Bio (Emily Hoelzli). Currently at Imprint Labs, LLC.

## Abstract

Using mRNA as a therapeutic has received enormous attention in the last few years, but instability of the molecule remains a hurdle to achieving long-lasting therapeutic levels of protein expression. In this study, we describe our approach for designing stable mRNA molecules by combining machine learning-driven sequence design with high-throughput experimental assays. We developed a high-throughput massively parallel reporter assay (MPRA) that, in a single experiment, measures the half-life of tens of thousands of unique mRNA sequences containing designed 3’ UTRs. Over multiple design-build-test iterations, we have accumulated mRNA stability measurements for 180,000 unique genomic and synthetic 3’ UTRs, representing the largest such dataset of sequences. We trained highly-accurate machine learning models to map from 3’ UTR sequence to mRNA stability, and used them to guide the design of synthetic 3’ UTRs that increase mRNA stability in cell lines. Finally, we validated the function of several ML-designed 3’ UTRs in mouse models, resulting in up to 2-fold more protein production over time and 30–100-fold higher protein output at later time points compared to a commonly used benchmark. These results highlight the potential of ML-driven sequence design for mRNA therapeutics.

## 1 Introduction

Messenger RNA (mRNA) has emerged as a transformative modality in the development of therapeutics and vaccines. Simple encoding of the therapeutic protein payload allows for rapid, modular design of novel drugs or vaccines. However, while small amounts of produced antigen are sufficient to elicit robust immune responses for vaccinations (e.g., Covid-19 vaccine), the relatively short half-life of mRNA is a significant hurdle to the production of the large quantities of proteins that are required for therapeutic applications (e.g., protein replacement therapies).

The 3’ untranslated region (UTR) of mRNA plays a critical role in regulating gene expression by influencing mRNA stability, localization, and translation efficiency [1]. Given that the protein-coding region of an mRNA is highly constrained for a given therapeutic, the design of the 3’ UTR sequence offers significant opportunities for enhancing the stability and efficacy of therapeutic mRNAs [2]. While codon usage and 5’ UTRs are routinely optimized to maximize protein production, including with ML-based methods [3, 4, 5, 6, 7, 8, 9, 10, 11, 12], attempts to use ML to design 3’ UTRs for enhanced stability or translation efficiency are in their infancy, even considering successful predictive modeling of various aspects of the 3’ UTR [13].

In this study, we designed high-stability synthetic 3’ UTRs by combining high-throughput data generation with ML-driven design. We developed a high throughput assay to measure the mRNA stability of thousands of 3’ UTRs in parallel using a massively parallel reporter assay (MPRA) framework, and we ultimately used this assay to curate the most extensive corpus of synthetic 3’ UTR sequences and corresponding stability measurements for 180,000 unique 3’ UTRs. We trained ML models on genomic and synthetic 3’ UTR stability measurements, and we found that our models were highly performant on test sets distinct from the training data. We explored the combination of ML model architectures with various design methods, exploring trade-offs between exploration and exploitation, and we were able to engineer synthetic 3’ UTRs with greater *in vitro* stability than observed in the training data. We further demonstrated the translatability of ML designs through validation of three high stability 3’ UTRs in an animal model. Following systemic injection of luciferase mRNA-LNP, whole-body bioluminescence imaging over a two-week period confirmed that our designed 3’ UTRs had longer half-lives than the control UTR, and by four days post-injection, our 3’ UTRs were driving up to 30–100-fold higher protein output. Notably, these 3’ UTRs demonstrated successful translation across different species, cell types, delivery modalities, and associated coding sequences (GFP and luciferase). We believe that our sequence discovery platform has the potential to enhance a wide range of mRNA therapeutics.

## 2 Highly reproducible Massively Parallel Reporter Assays construct training data for learning the function of 3’ UTR sequence to mRNA stability

We first developed and performed multiple Massively Parallel Reporter Assays (MPRAs) to measure the stability of mRNA *in vitro* in a high throughput assay while varying the 3’ UTR of the mRNA (Figure 1 a). mRNA stability is measured in an MPRA by linking 3’ UTRs to a reporter gene (primarily 2deGFP), where relative amounts of each 3’ UTR are quantified with high throughput sequencing at four time points. Relative counts across time points are then used to estimate the decay rate, or half-life, for each 3’ UTR (See A.2.6). Figure 1 b demonstrates degradation curves for an exemplary assay of 14,460 3’ UTRs across four time points. Fluorescence-activated Cell Sorting (FACS) at 6hr and 24hr time points for four 3’ UTRs confirmed variation in protein expression of a subset of 3’ UTRs, with notable differences in expression at the 24hr time point (Figure 1 c). In general, we observed remarkably high repeatability across experiments when comparing half-lives of control 3’ UTRs across experiments with similar uridine modifications (Figure A1 c).

**Figure 1:**
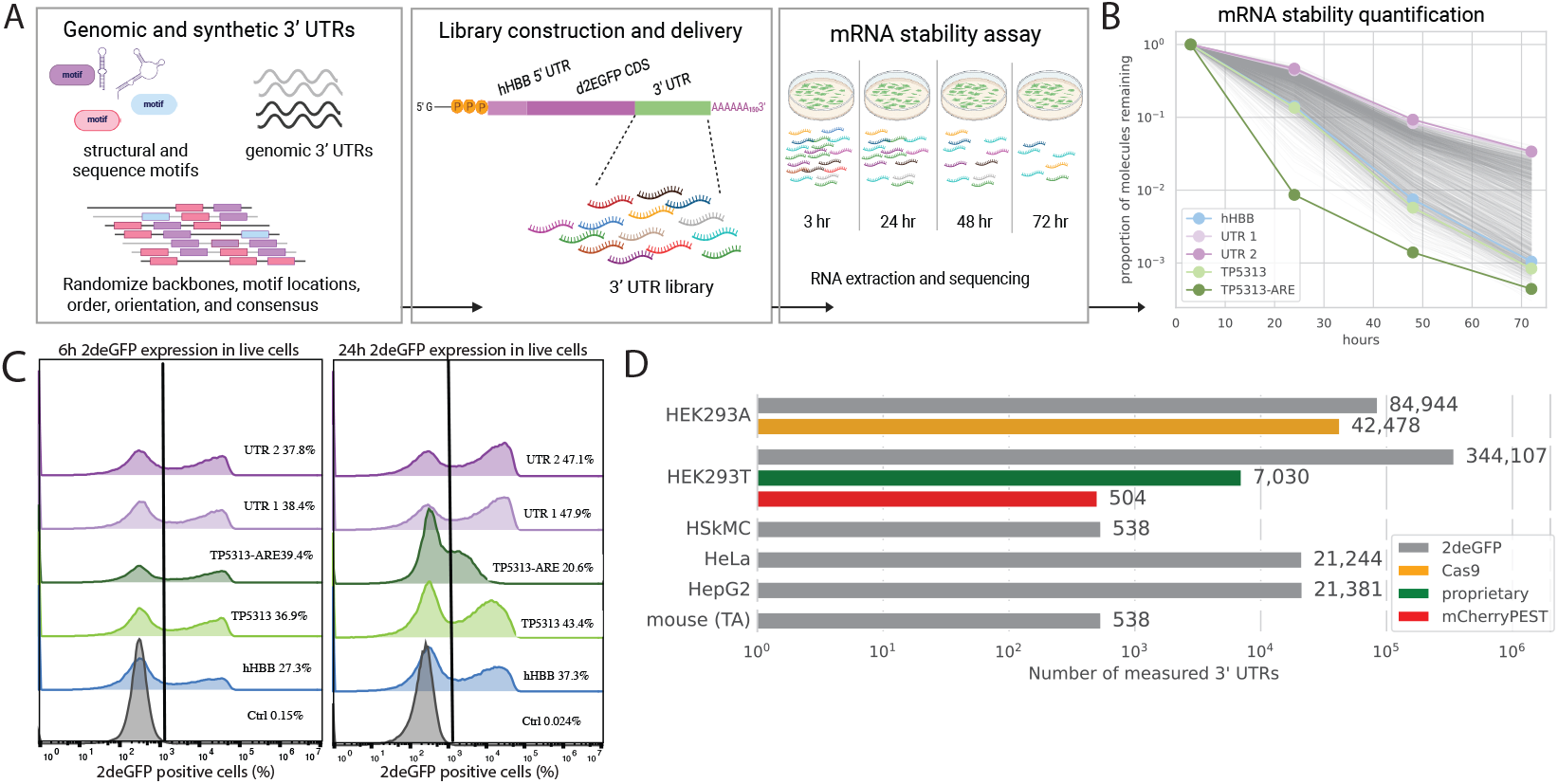
High throughput measurements of mRNA stability for ∼180,000 unique mRNAs with a variable 3’ UTR were collected using massively parallel reporter assays (MPRA). (a) Experimental overview for measuring stability of mRNAs with variable 3’ UTRs *in vitro*. 3’ UTRs were inserted into a plasmid backbone, mRNAs were transcribed, capped, and polyadenylated, and finally transfected into cell lines. Cells were incubated over 4 time points, after which mRNA was quantified for each 3’ UTR candidate with next generation sequencing (NGS). 3’ UTRs were collected from genomic sourcing or designed synthetically as described in Methods A.1.2. (b) Degradation curves for an exemplary MPRA containing 14,460 unique 3’ UTRs. Mean RNA counts from three replicates were collected from 4 time points (3hr, 24hr, 48hr, and 72hr). Counts are normalized to 3 hr time point. (c) Fluorescence-Activated Cell Sorting (FACS) at 6 hr and 24 hr time points for five cell populations each containing a gene cassette consisting of the hHBB 5’ UTR, a 2deGFP coding sequence, and one of four 3’ UTRs as shown. Percentages indicate the percent of GFP positive cells at each time point. The four 3’ UTRs are derived from the human hemoglobin subunit beta (hHBB) gene, the human TP5313 gene and a similar cassette including an adenylate-uridylate-rich element (ARE), and two synthetic 3’ UTRs. (d) 3’ UTRs were measured in five cell lines (“*in vitro*”) and in a mouse model (“*in vivo*”, Tibialis Anterior (TA)). 3’ UTRs were measured with the hHBB 5’ UTR and four different coding sequences.

We measured half-lives for an initial set of 16,586 “natural” genomic 3’ UTRs and 25,893 synthetic 3’ UTRs (see A.1.1 and A.1.2). We trained ML models on this initial dataset and performed three iterations of ML-driven design of 3’ UTRs. In total, we generated stability measurements for 180,000 unique 3’ UTRs. However, we collected additional measurements while varying the cellular condition, 5’ UTR, and CDS, where each assay contained shared 3’ UTRs to support comparison across conditions and normalization (Figure 1 d). To our knowledge, this is the largest dataset that measures the effect of 3’ UTR design on mRNA stability.

## 3 ML-driven design of 3’ UTRs demonstrates increased mRNA stability over three iterations of design

We trained supervised models using our initial dataset of 42,479 genomic and synthetic 3’ UTRs to learn the function of 3’ UTR sequence to mRNA stability (Methods A.6, further discussed in Section 4). We developed and tested seven design methods and compared their ability to produce novel, diverse, and stable 3’ UTRs that exceeded stability measurements in the training dataset. Specifically, we tested a genetic algorithm ([14]), simulated annealing, model driven mutagenesis, an implementation of activation maximization (SeqProp [15]), and implementations of DbAS [16] and CbAS [17] (Figure 2 a, Methods A.10). As a baseline, we randomly inserted known structural and canonical sequence motifs into backbones and used a supervised model to filter 3’ UTRs predicted to be stable (referred to as *Synthetic (ML selected)*, Methods A.10.7). While all design methods primarily used a supervised LSTM architecture (Methods A.6.1) to predict the stability of 3’ UTRs (Figure 2 a), a subset of these methods used an unsupervised masked language model or a variational autoencoder (Methods A.8, A.7). Both unsupervised models were trained on 3’ UTR sequences from UTRdb [18].

**Figure 2:**
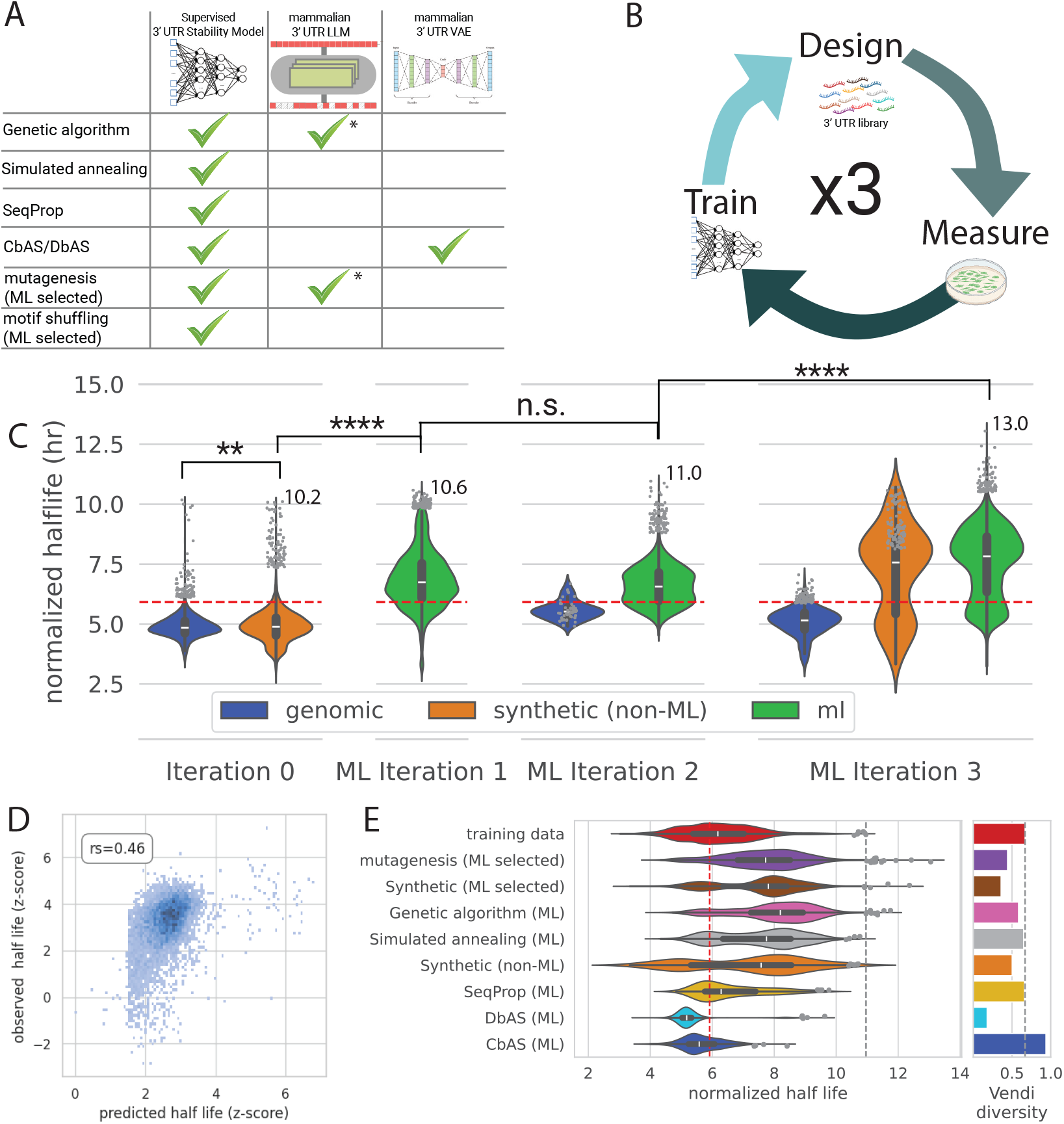
ML-driven design of 3’ UTRs. (a) Methods that were used to design 3’ UTRs. Seven methods used a supervised model, large language model (LLM) and/or variational autoencoder (VAE) trained on 3’ UTRs from 125 mammalian species. **\***Large language models (LLMs) were used in a subset of genetic algorithms and mutagenesis designs to test different methods for selecting mutations. (b) Schematic of design cycle. We ran three iterations of ML-driven design. In each iteration, *in silico* designs were generated using methods shown in Figure a. Designs were validated *in vitro* in an mRNA stability MPRA, and models were re-trained with additional data after each round. (c) Comparison of measured half-lives of mRNA containing variable 3’ UTR regions of genomic and synthetic designs across three design iterations. Half-lives were normalized across assays using shared carryovers. Horizontal red line indicates normalized half-life for hHBB 3’ UTR benchmark. All mRNA shown includes 2deGFP reporter gene and hHBB 5’ UTR. (d) Correlation between predicted and measured stability (z-score normalized half-lives) of mRNA that includes 3’ UTRs designed using the model shown. (e) Comparison of measured half-lives of mRNA containing ML-designed 3’ UTRs, separated by design method. Vertical red line indicates normalized half-life for hHBB 3’ UTR benchmark. All mRNA shown includes 2deGFP reporter gene and hHBB 5’ UTR. (right) Vendi score measures sequence diversity for each ML design method. Vendi score is calculated using normalized kmer frequencies for each 3’ UTR.

We performed three iterations of ML design, where each iteration included a cycle of ML-driven design, experimental validation through our stability MPRA, and re-training supervised models on new measurements (Figure 2 b). We observed significant increases in the stability of sequences designed in ML iterations 1 and 3, when compared to the previous iteration (Figure 2 c). Notably, all three iterations of ML-driven design propose 3’ UTRs that have improved stability over the commonly used 3’ UTR benchmark from hHBB [19, 20, 21]. Stability predictions for designed UTRs from an exemplary supervised model had reasonable correlation to measurements collected in the subsequent build iteration (rs = 0.46, Figure 2 d). This correlation demonstrates that on average our models can reliably design 3’ UTRs that translate *in vitro*. Outliers to this trend in Figure 2 d demonstrate a small number of 3’ UTRs whose predicted stability exceeds that of measured stability (predicted z-score normalized half-life > 4, Figure 2 d). In this particular case, all outliers were designed with the same design method (mutagenesis selected with predictive model, see Methods A.10.7). These designs highlight regions of the sequence space where our models were unable to reliably predict.

While stability of designed sequences significantly increased in the first and third ML design iterations, stability from the first to second iteration did not significantly increase (p-value = 1.0, Figure 2 c). Additionally, PCA of ML designs, calculated from supervised LSTM model embeddings, suggest that while designs from the third ML iteration are uniquely distinct from synthetic and genomic sequences (Figure 4 c), designed UTRs from ML iteration 2 occupy a qualitatively similar area in the latent space as designs from iteration 1 (Figure4 c). These results suggest that while the first and third iterations of ML design succeeded in pushing stability significantly higher than past iterations while exploring unique sequence spaces, designs from iteration 2 did not achieve these milestones.

**Figure 3:**
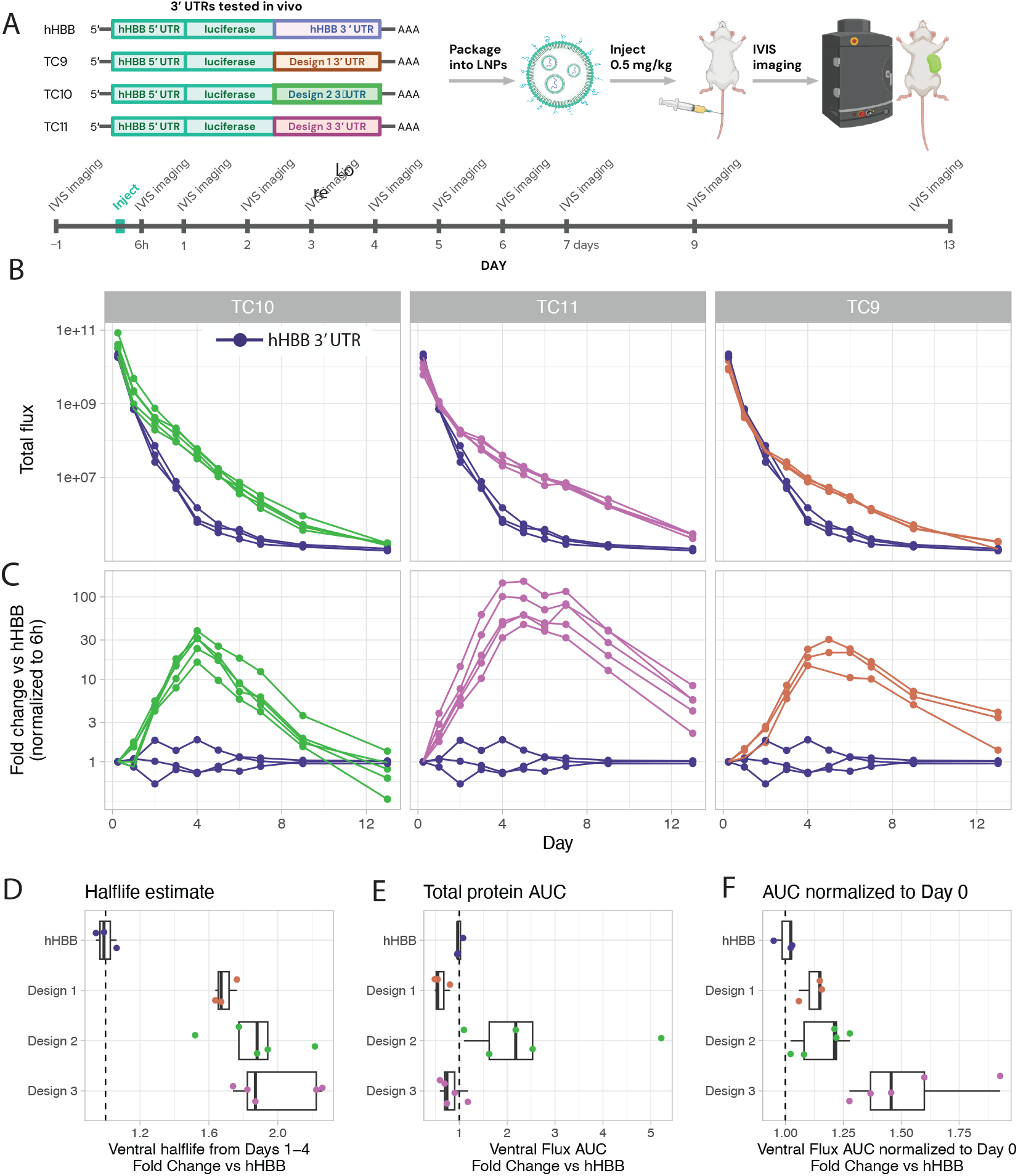
*in vivo* mouse validation of 3’ UTRs (a) mRNAs containing three 3’ UTR candidates and the hHBB 3’ UTR benchmark were LNP-formulated for *in vivo* validation. *in vivo* Imaging System (IVIS) imaging was collected over 13 days. (b) Total flux over 13 days for 3’ UTR candidates. Fold changes were normalized to 6 hour IVIS imaging time point. (c) Fold change over hHBB 3’ UTR benchmark over 13 days for 3’ UTR candidates. Fold changes were normalized to 6 hour IVIS imaging timepoint. (d) Half-life estimates for hHBB and three ML designed sequences based on Days 1 to 4. (e) Total protein estimates based on area under the curve (AUC). calculation. (f) Total protein estimate based on area under the curve (AUC), normalized to Day 0 to account for potential differences in formulation or delivery quality.

**Figure 4:**
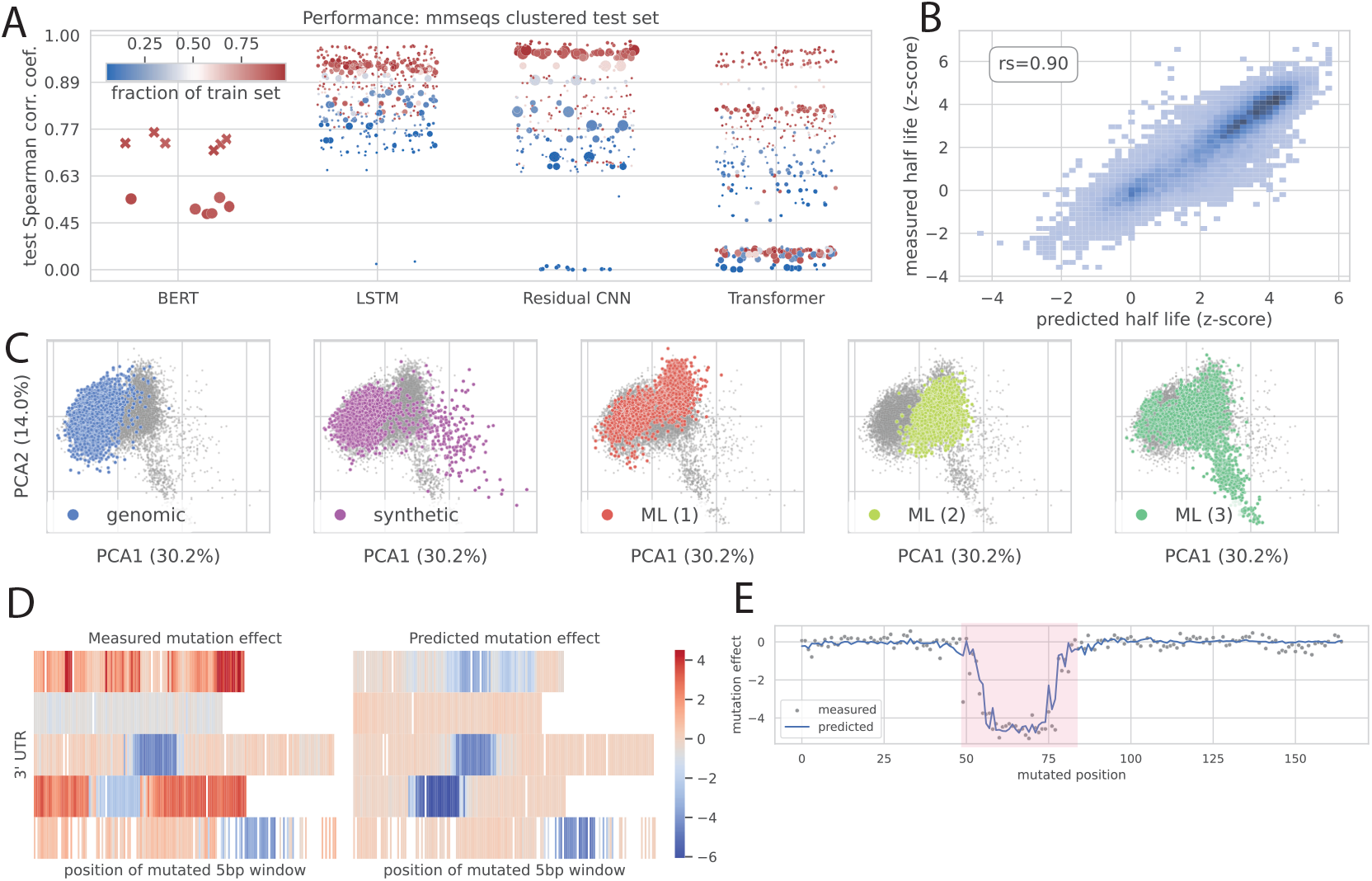
Model architectures vary in their ability to predict effect of 3’ UTR sequence on mRNA stability. (a) Comparison of Spearman correlation on held out test set across four distinct supervised model architectures (detailed in Section A.6). BERT models were either pre-trained on mammalian 3’ UTRs before fine tuning on stability data (marked with X) or fine tuned from a random initialization of parameters (marked with circles). y-axis: non-linearly transformed to differentiate models with high test Spearman correlation coefficients. Size of each datapoint represents the number of parameters for each model, which range from 1,345 to 43,896,577. (b) Comparison between predicted and observed stability measurements in a held out test set for best performing LSTM model (about 84k parameters). Spearman’s rank correlation coefficient = 0.90. (c) PCA of LSTM model embeddings for a subset of 3’ UTRs (2000 UTR’s from each category: genomic, synthetic, and ML designed sequences from 3 iterations). Embeddings were derived from the output of the first convolutional layer before LSTM layers. UTRs are colored by whether they are genomic, synthetic (human-designed and selected), or ML designed. (d) (Left) measured and (right) predicted effect sizes for 3’ UTRs that underwent window mutagenesis (5bp sliding window of random mutations). (e) Predicted and measured effect sizes for window mutagenesis for an exemplary 3’ UTR. Effect sizes are calculated as the predicted (or measured) z-score transformed half-life of a mutant with five consecutive nucleotides mutated minus the predicted (or measured) z-score transformed half-life of the unmutated 3’ UTR.

### 3.1 ML-driven design methods trade off exploitation of measured mRNA stability with sequence diversity in a single iteration of design

We next evaluated methods on their ability to design stable and diverse 3’ UTRs. As a basic assessment, we first confirmed that synthetic 3’ UTRs selected with a supervised stability model outperformed random selection significantly outperformed random selection (Supplementary Figure A2 b). Upon confirmation of the model’s ability to select stable sequences, we next tested and compared additional design methods shown in Figure 2 e. Surprisingly, designs with the highest stability were generated by performing *in silico* mutagenesis on stable UTRs and selecting mutations that were predicted to increase stability via a supervised model (Methods A.10.7). This design method is greedy in nature, resulting in both highly stable designs and and a subset of designs that did not translate experimentally (Figure 2 d).

We explored various methods for selecting mutations for *in silico* mutagenesis and our genetic algorithm. Methods included random selection of mutations, selection with a supervsied model, and selection with a 3’ UTR large language model (LLM) trained on genomic mammalian 3’ UTRs (Methods A.10.7,A.10.8). Selection of mutations during *in silico* mutagenesis using a supervised model significantly outperformed selection of mutations using likelihoods from our 3’ UTR LLM (Methods A.10.7,A.10.8, Supplementary Figure A2 a). Similarly, selection of mutations in a genetic algorithm using a supervised model resulted in sequences with highest stability, when compared to random selection and likelihood based selection from our 3’ UTR LLM (Methods A.10.1, Supplementary Figure A2 c). Given that likelihoods of the genomic 3’ UTR LLM do not take stability into account, it is unsurprising that this method would produce less stable designs. However, in these approaches LLM’s introduce mutations based on their likelihood of occurrence in genomic sequences, potentially producing designs that are more similar to genomic 3’ UTRs than purely synthetic designs.

Because unsupervised models were used for a subset of designs (refer to Figure 2 a), we sought to understand the effect of training data on quality of designs proposed by unsupervised models. To this end, we designed UTRs using an implementation of CbAS and DbAS, which implements a variational autoencoder (VAE) to sample from the sequence space. We trained two VAEs, one of which was trained on 125 mammalian species (mammalian VAE) and another which was trained on 550 species not limited to mammalian species (multi-species VAE, Methods A.7, A.5). Supplementary Figure A2 d demonstrates that although the multi-species VAE was trained on significantly more data, the mammalian VAE produces designs with higher overall stability. Considering that our designs were validated in mammalian *in vitro* and *in vivo* systems, these results emphasize the importance of aligning training data to the sequence distribution most relevant to the validation system.

To assess diversity for all design methods, we compute the Vendi score [22], a metric to summarize diversity, computed from the relative frequencies of 8-mers in each UTR (Figure 2 e). While methods such as DbAS design UTRs with relative low mean stability and diversity, other methods such as simulated annealing demonstrate both higher mean stability in addition to higher diversity. Simulated annealing, Seqprop, and CBAS demonstrate similar or increased diversity to that the training data, which is required to promote exploration and avoid quick convergence to local optima in subsequent design iterations [17, 23, 24].

Methods discussed so far do not encourage exploration of the sequence space in areas that the model cannot predict in. This drawback may inhibit discovery of novel sequences in next design iterations. Active learning aims to improve model generalization over time by selecting data points that are informative to the model. To test this, we designed 15 million synthetic sequences and used three active learning methods, including core-sets [25], margin-sampling [26], and cluster-margin [27], to select 2,500 synthetic sequences per method and assess how each method improved model performance. Surprisingly, including sequences selected using active learning did not improve performance on a held out test set when compared to randomly sampled sequences (Supplementary Figure A2 e). Cluster-margin [27] and margin sampling [26] require estimates of model uncertainty, in which we used standard deviations from an ensemble of three models (See Methods A.11). As we discuss in Section 5, ensembling is poorly calibrated for estimating uncertainty [28] which may negatively affect results. Additionally, active learning is highly reliant on the initial set of data points [29]: if these data are too similar to the training data, active learning is unlikely to provide performance gains. These inherent limitations of active learning methods make it difficult to assess its value in isolation from the choice of initial data points and the method for quantifying uncertainty.

### 3.2 ML designed 3’ UTRs translate across contexts and increase protein expression *in vivo*

One of the major challenges in therapeutic development is utilizing accurate model systems that effectively predict drug success in human patients. To this end, we collected additional measurements of our 3’ UTR libraries while varying the cell line, 5’ UTR, CDS, and mRNA nucleotides (Figure 1 d) to determine the effect of cell condition and full mRNA sequence on mRNA stability (Supplementary Figures A1 d,e,f). While it is expected that mRNA stability will vary between cell types due to differential expression of RNA-binding proteins and miRNAs [30, 31, 32], we observed moderate variation in stability between the HepG2, HeLa, Primary Human Skeletal Muscle Cells (HSkMC), and HEK293T cellular conditions (Spearman’s rank correlation coefficient ranges from 0.54 to 0.70, Supplementary Figure A1 b). We observed similar variation across assays when swapping out the CDS and the 5’ UTR (Spearman’s rank of 0.90 and 0.66, Supplementary Figures A1 e, A1 f), which have been shown to impact mRNA stability through modification of translation initiation, translation speed, and ribosome pausing [33, 34, 35]. Finally, our original assays were performed with unmodified nucleotides, but modified nucleotides are often used in mRNA therapeutics due to their increased translational capacity and reduced immunogenic effects [36, 37]. We repeated our assays with N1-methylpseudouridine instead of unmodified uridine and found a reasonable difference in 3’ UTR behavior (Spearman’s rank correlation coefficients -0.1 to 0.9; Supplementary Figure A1 c). Interestingly, we saw that low and moderate stability 3’ UTRs from the original assay were indistinguishable from each other in the modified nucleotide assay, while the highest stability 3’ UTRs were shared between both assays (Supplementary Figure A1 d).

As a final test of translation, we selected three ML-driven designed 3’ UTRs for *in vivo* validation of protein expression. These 3’ UTRs were selected for their high mRNA stability *in vitro* and for diversity in sequence features. Selected 3’ UTR sequences and the hHBB 3’ UTR benchmark were formulated as mRNA-LNPs with a hHBB 5’ UTR and luciferase coding sequence (Figure 3 a). mRNA-LNPs were injected systemically into mice, and whole-body bioluminescence quantification was performed at multiple time points over 13 days (Figure 3 a).

Whole-body IVIS imaging results are shown in Supplementary Figure A3 a, with protein activity measurements in total flux (photons/second; Figure 3 b) and relative flux normalized to Day 0 measurements versus hHBB 3’ UTR animals (Figure 3 c). By day seven following injection, all three ML-driven designs showed higher protein activity than the hHBB 3’ UTR benchmark, with differences reaching 100-fold for Design 3 (Figure 3 c). Notably, luciferase protein has a relatively short half-life of approximately three hours [38]. Due to the luciferase protein instability, measured protein activity is expected to correlate highly with current mRNA abundance. We estimated mRNA half-life per animal based on the protein activity from Days 1 to 4, before any animals reached our lower limit of detection (*<* 5 × 10^5^ total flux), and found that ML-driven designs showed up to 2-fold increases in half-life versus hHBB benchmark (Figure 3 d).

To differentiate total protein expression across UTRs, we calculated total protein expression using absolute area under the curve. However, day 0 measurements varied greatly between 3’ UTRs and in some cases between animals injected with the same 3’ UTR – four animals were removed from further analyses due to outlier flux measurements at day 0. We calculated total protein expression using absolute area under the curve calculations (Figure 3 e) and using relative area under the curve with flux measurements normalized to day 0 due to previous described variation at day 0 (Figure 3 f). Protein expression estimates vary between calculation methods for AUC, though Design 2 shows higher protein expression than the benchmark 3’ UTR with both calculations (Figure 3 f).

## 4 Architectures vary in their ability to predict effect of 3’ UTR sequence on mRNA stability

During design iterations, we evaluated the predictive accuracy of various model architectures on 3’ UTR half-life measurements in HEK293T to determine whether different ML model architectures exhibited inductive biases that either promoted or inhibited their ability to learn the effect of 3’UTR sequence on mRNA stability. To this end, we evaluated four model architectures on a held out test set of 3’ UTRs and corresponding half-lives. Architectures included three supervised models (transformer, long short term memory (LSTM), and convolutional neural network, Methods A.6), as well as a semi-supervised BERT model [39] that was pre-trained on 3’ UTRs from 125 mammalian species selected from UTRdb [18].

### Training data

To ensure that models were evaluated on their ability to generalize to distinct 3’ UTRs, we curated a test set of 3’ UTRs by clustering 3’ UTRs using mmseqs2 [40] and holding out a subset of entire clusters. Because we were ultimately interested in applying these models to design 3’ UTRs that increase mRNA stability over what was observed in the training data, we additionally included all clusters that contained the most stable 3’ UTRs to assess each model’s ability to predict half-lives that were greater than what was observed during training. Ultimately this yielded a dataset of 113,414 unique 3’ UTRs, with 27,311 clusters in the training set and 135 clusters held out for evaluation. The average sequence length of UTRs was approximately 164 nt.

We trained a set of models while varying the number of model parameters, fraction of mmseqs2 clusters used for training, and model architecture. During training, subsets of clusters were randomly selected to curate a training set, resulting in models that were trained on 5% to 96% of the available data. While testing different model architectures allowed us to evaluate the differences in inductive biases driven by architecture choice, varying the number of model parameters and the fraction of data used during training allowed us to observe data and model parameter scaling laws empirically.

Figure 4a demonstrates the performance of all models on the described test set. While top convolutional neural network models performed slightly better than top LSTM models (n = 40, t-test FDR = .004), transformers performed worse than both LSTMs and residual convolutional models (FDR = 2.2e-19), with a large variation in predictive accuracy that was largely driven by the number of parameters (Supplementary Figure A4c). Interestingly, all three supervised architectures reached maximum Spearman correlation on the held out test set around 10e6 parameters. For larger models, we observed degradation in performance with more than 10e6 parameters (Supplementary Figure A4a-c). Unlike the LSTM and CNN, transformer performance dramatically decreased with larger architectures. Additionally, while all models demonstrate improvement upon the addition of more training data when parameter number is held constant (Supplementary Figure A4a-c), transformers demonstrate larger variation in accuracy when training data size is varied (Supplementary Figure A4c). We hypothesize that this decrease in performance is likely due to overfitting on the train set.

Figure 4a additionally shows that the performance of a BERT model fine tuned on our stability data significantly increases when pre-training on mammalian 3’ UTRs, when compared to a randomly initialized Bert model (FDR = 1.43e-8). However, the fine tuned BERT architecture is not as performant as supervised architectures, despite its drastically greater number of parameters (Figure 4a). These results demonstrate that smaller supervised models can outperform larger foundational models when learning from functional datasets, and should be first evaluated before defaulting to fine tuning a foundational model.

Due to the parameter efficiency and performance of the LSTM and its robustness to training dataset size (Supplementary Figure A4 a), we primarily utilized an LSTM architecture as discussed in Section A.6.1. While Figure 4b demonstrates high performance on the test set (Spearman correlation coefficient = 0.90), principal component analysis (PCA) of model embeddings for held out 3’ UTRs visually demonstrate that the model differentiates between genomic sequences, human designed synthetic sequences, and ML designed synthetic sequences (Figure 4c). Interestingly, Figure 4c demonstrates that in PCA coordinates of embeddings for the sequences from the third iteration of ML design sequences occupy a distinct space, when compared to genomic, synthetic, and ML designed sequences from the first two iterations of ML design. This observation qualitatively demonstrates feedback covariate shift [33] between ML designed and the original training dataset over time.

### 4.1 *in vitro* and *in silico* validation of models identify motifs that affect mRNA stability

While our models were performant on a held out test set, we next sought to determine whether models were learning regulatory grammar of 3’ UTRs that affect mRNA stability. To test the ability of our model to learn motifs that affected mRNA stability *in vitro*, we held out an additional portion of our training dataset that consisted of randomly mutated windows of 5 consecutive base pairs at each position in a set of 3’ UTRs, in addition to the original test set described. Figure 4d demonstrates 5 instances of measured and predicted effects of window mutagenesis on 3’ UTRs, where predictions mimic patterns of measured window mutagenesis for four of five sequences. Indeed, we observed that one of the regions of a 3’ UTR that drastically decreased mRNA stability when mutated (Figure 4e) included a motif that was used to construct human designed synthetic 3’ UTRs A.1.2. Running TF-MoDISco [34] on model attributions confirmed that our model was learning this motif as a significant motifs that determined mRNA stability (data not shown).

While *in vitro* measurements can anecdotally validate small sets of motifs in the 3’ UTR that affect mRNA stability, we wanted to comprehensively evaluate which motifs that were originally mined for training data (See A.1.2) were significantly affecting mRNA stability when inserted into the 3’ UTR. To test the effect of each motif on mRNA stability, we randomly inserted each motif into various backbones and performed a t-test under the null hypothesis that a given motif did not significantly alter mRNA stability. We found that of the 212 motifs originally mined for training data generation, 177 were predicted to significantly increase mRNA stability when present at least once in a 3’ UTR (FDR < 0.05, Supplementary Figure A4e). Additionally, when using two separate model architectures to compute statistics, both models agreed on both the effect size and significance of motifs (Supplementary Figure A4e).

To determine whether any motifs had synergistic effects, we generated a synthetic set of sequences that inserted different combinations of motifs at different distances into randomly selected 3’ UTRs. We then fit a ridge regression model using our model’s predictions of the synthetic 3’ UTRs to determine whether any motifs had synergistic or antagonist effects. Of the 212 motifs inserted, we observed no synergistic effect between motifs, while we observed only four antagonistic interactions (FDR < 0.05, Supplementary Figure A4f). As many of these motifs were relatively long when compared to the length of the 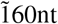 3’ UTR (covering 16% of the 3’ UTR on average), we hypothesize that structures formed within these motifs may interfere with local structure of additional motifs and therefore interfere with mRNA stability [35, 36, 41]. This lack of observed synergistic motif effects differs from work that models enhancer and promoter activity [42]. These differences in underlying regulatory grammar may therefore require different considerations when constructing both training sets and selecting model architecture for RNA design.

## 5 Discussion

We present our approach for designing stable mRNA molecules by combining machine learning models, ML-driven sequence design methods, and high-throughput experimental assays. We demonstrated how our approach to designing 3’ UTRs successfully increased mRNA stability in cell lines, and that those designs translated to mouse models. The iterative nature of our platform (a design-build-test loop) also produced mRNA stability measurements of 180,000 unique 3’ UTRs, representing the largest such dataset to our knowledge.

Although we observed significant improvement in two iterations of ML-driven design, increases in the maximum stability in each iteration were low. These results may suggest a trade-off between taking larger steps out-of-distribution and reliably generalizing to new sequences. While we have investigated various methods to determine whether designs will translate to model systems, we found common methods such as ensembling are poorly calibrated towards model error under a distribution shift, similar to findings in [28]. Heuristics such as a designed sequence’s distance from the training data (computed from both k-mer distance and model embedding distance) did correlate with predictive error (Pearson = 0.44). However, predictive error correlated most strongly with model predictions (Pearson = 0.9), suggesting that simply constraining maximum predictions of designs is sufficient to ensure reliable predictions. While guiding designs by constraining maximum predictions may improve success of designs in a model system, it will not help us reach our ultimate goal of designing sequences with much greater stability than what was observed in the training data.

A large challenge in therapeutic development is using accurate model systems that translate to prolonged protein expression in human patients. To this end, we performed early experiments across conditions, including variations to cell type, 5’ UTR, and coding sequence, allowing us to confirm certain sequence features that translate between contexts. The translation of our designed UTRs’ effects on mRNA stability across contexts was ultimately confirmed by our *in vivo* study, which changed species, cell type, coding sequence, and packaging while maintaining increased mRNA stability. However, the relationship between mRNA stability and protein expression is less straightforward [20]. Early experiments confirmed that our top stable 3’ UTRs did show increased protein expression at 24 hours (Figure 1c), but our *in vivo* results found that absolute protein production AUC was reduced for two of our 3’ UTR candidates versus the benchmark (Figure 3e), though mRNA half-life was estimated to increase (Figure 3d). Though we cannot rule out differences in mRNA-LNP quality or delivery between UTRs in our *in vivo* experiment, our results should serve as a caution to the community to consider the complete context translation when optimizing sequence features on the path to therapeutic development.

## Acknowledgments and Disclosure of Funding

This research was funded by Ginkgo Bioworks.

We would like to thank Jennifer Listgarten, Tal Ashuach, and Anshul Kundaje for their helpful feedback during their time with Patch Biosciences. We would like to thank Lood van Niekerk, Seth Ritter, and Justin Gardin for their helpful comments and contributions to the paper.

---

**Figure A1:**
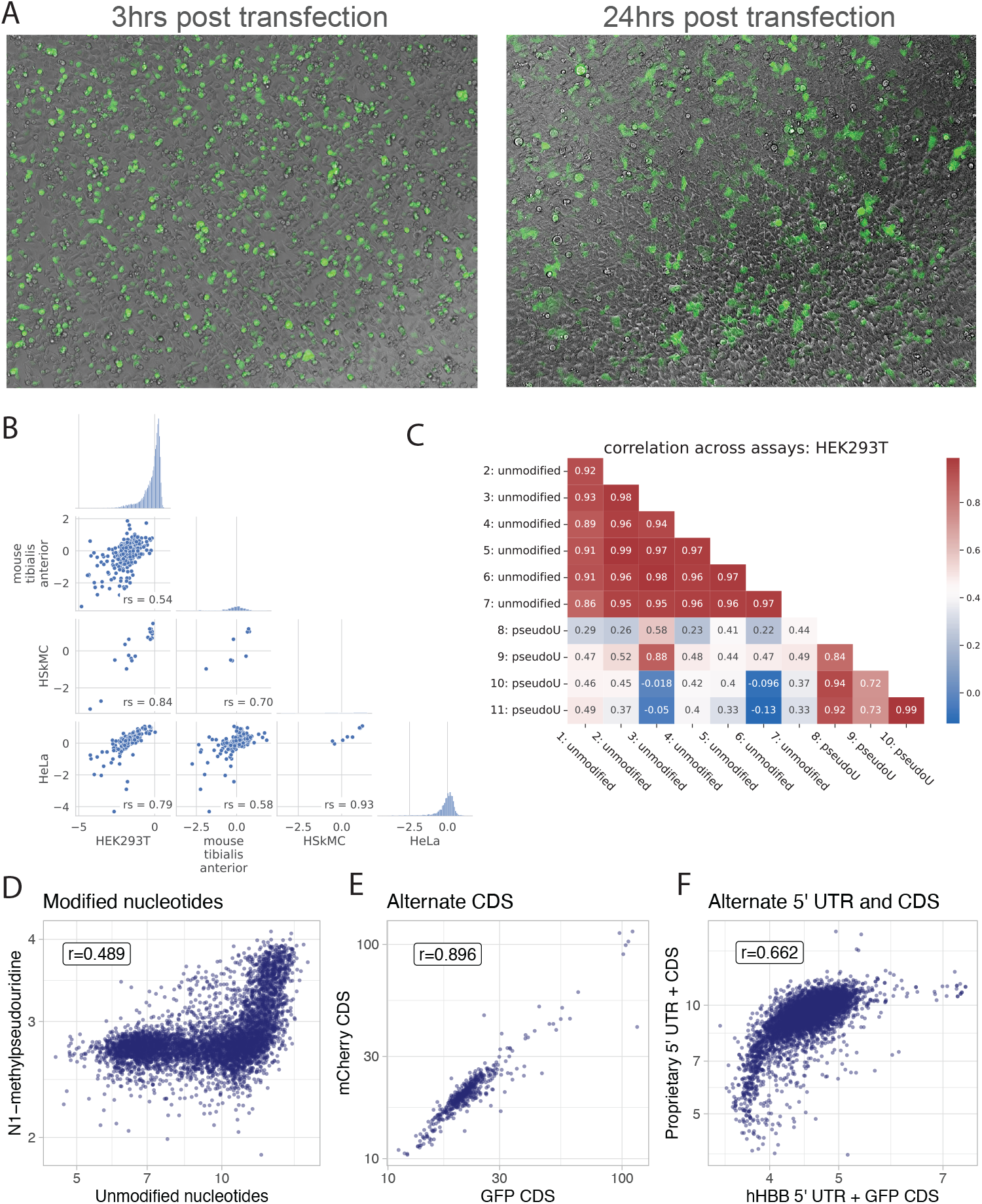
Correlation across MPRAs (a) GFP expression in HEK293T cells 3hrs and 24hrs post mRNA transfection. GFP expression of HEK293 cells was imaged using Evos fluorescence microscope under 10x magnification. (b) Spearman’s rank correlation coefficient: measured half-lives between MPRA assays with shared 3’ UTRs in *in vitro* conditions HEK293T, Primary Human Skeletal Muscle Cells (HSkMC), HeLa, and *in vivo* mouse tibialis anterior. (c) Spearman’s rank correlation coefficient of measured half-lives between MPRA assays with shared 3’ UTRs. Assays are labeled as unmodified uridine and N1-methyl pseudouridine. Correlations were calculated after removing low-quality measurements. (d) Spearman’s rank correlation coefficient and corresponding normalized half-lives RNA counts between two MPRAs with unmodified uridine and N1-methyl pseudouridine. (e) Spearman’s rank correlation coefficient and corresponding normalized half-lives RNA counts between two MPRAs with different coding sequences (CDS): 2deGFP and 2demCherry. (f) Spearman’s rank correlation coefficient and corresponding normalized half-lives RNA counts between two MPRAs with different coding sequences (CDS) and 5’ UTR.

**Figure A2:**
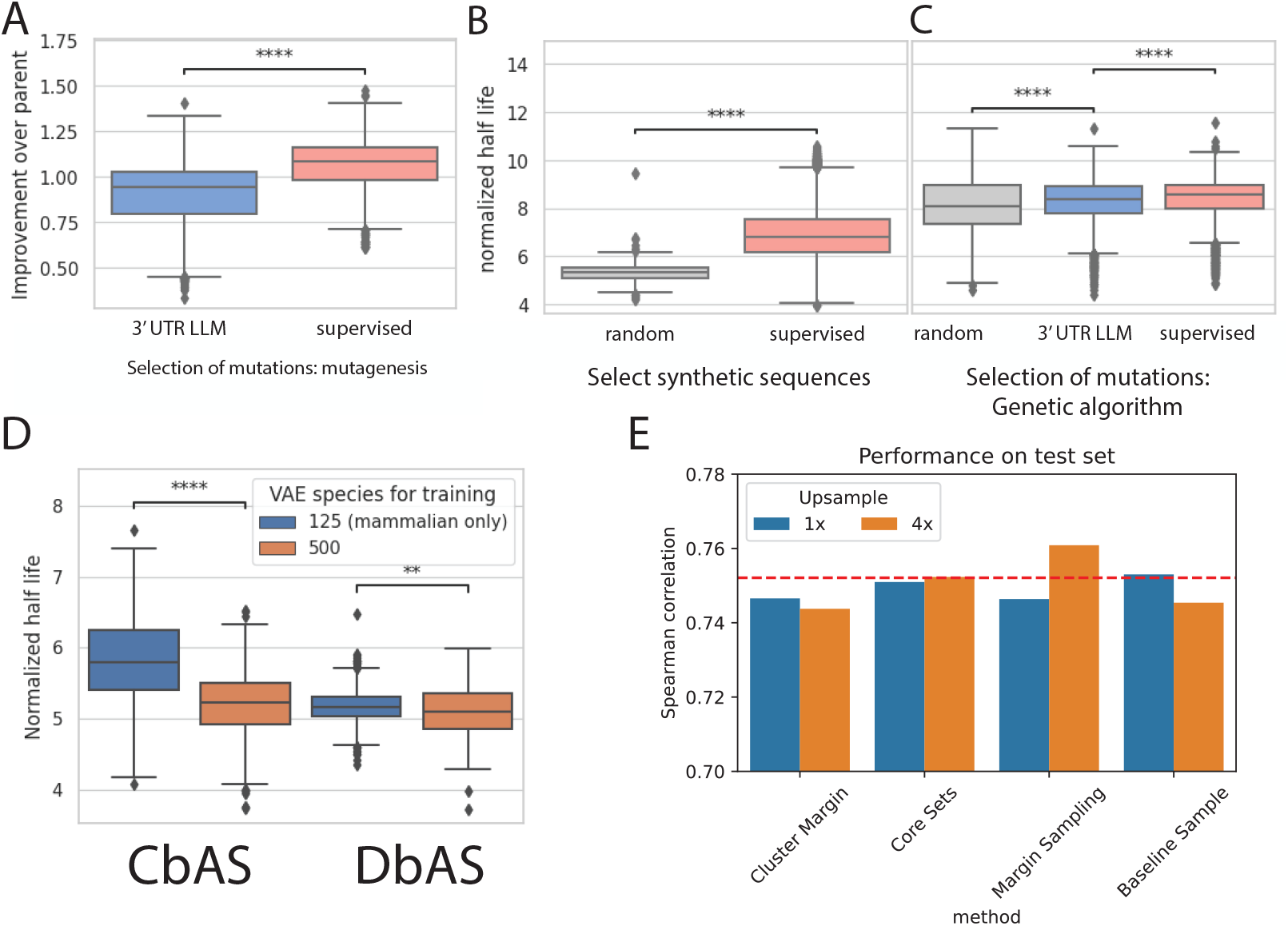
Comparison of alternative methods for ML-driven design (a) Improvement of half-lives for mutated sequences compared to original sequences. 1.0 indicates no improvement of mutated sequence over parent. Mutations were selected based on likelihoods from a large language model trained on 3’ UTRs from 125 mammalian species (3’ UTR LLM), or were selected based on predictions of mutated sequences using a convolutional neural network (supervised). (Mann-Whitney-Wilcoxon one sided, p-value = 4.761e-186) (b) Measured half-lives for synthetic sequences that were randomly selected or selected using an ML model based on predictions. (Mann-Whitney-Wilcoxon one sided test, p-value = 2.429e-129) (c) Measured half-lives for sequences designed with a genetic algorithm. During design mutations were either randomly selected (random), selected based on likelihoods from a large language model trained on 3’ UTRs from 125 mammalian species (3’ UTR LM), or were selected based on predictions of mutated sequences using a convolutional neural network (CNN). (Mann-Whitney-Wilcoxon one sided, p-value = 1.316e-07,9.436e-11) (d) Measured half-lives for sequences that were generated using CbAS and DbAS methods, which sample from variational autoencoders (VAE) during design. Two variational autoencoders were trained on 3’ UTRs and used to generate sequences. The first VAE trained 3’ UTRs from 125 mammalian species. The second VAE trained on 3’ UTRs from 500 species (including, but not limited to, mammalian species). (e) Comparison of active learning methods. Models were re-trained with 1,300 to 1,700 UTRs that were selected using Cluster-Margin, Core-Sets, or Margin Sampling active learning methods, as well as a baseline that randomly sampled from a set of synthetic 3’ UTRs, weighted by predicted activity. Models were trained with active learning selected UTRs in addition to 62,000 UTRs previously measured. Horizontal line indicates performance of model with no active learning sequences included in the training set (62,000 sequences). We upsampled sequences chosen by active learning during training to boost their representation in the training set (no upsampling (1x, blue), or upsampling active learning representations by 4 (4x, orange).

**Figure A3:**
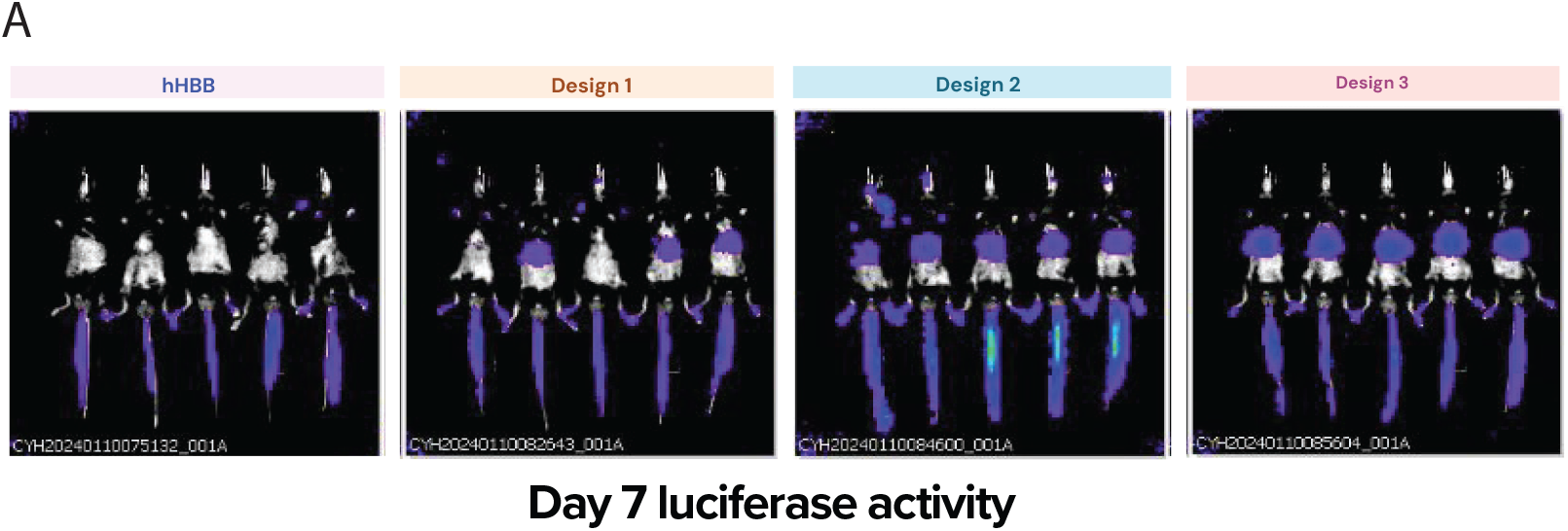
*in vivo* validation (a) Luciferase activity for three 3’ UTR candidates and the hHBB 3’ UTR benchmark at day 7. Four replicates per UTR.

## A Supplemental material

### A.1 3’ UTR sequence design

#### A.1.1 Selection of genomic and control 3’ UTRs for training data curation

Our initial 3’ UTR dataset was derived from genomic 3’ UTRs in highly expressed genes. We selected the top 1000 expressed genes in various cell lines based on data from the Cancer Cell Line Encyclopedia (CCLE) [43], and we created overlapping 170-nt tiles of the 3’ UTRs and their reverse complement sequences. These 37,971 sequences formed our initial genomic 3’ UTR dataset.

We also curated a list of 502 control 3’ UTRs to use for data quality control across assays. This dataset included 185 full length and tiled 3’ UTR sequences from previously published 3’ UTR screens [44, 18], 167 sequences with previously discovered activating and repressive 34mers [45] shuffled into randomer backbones, and 166 sequences that showed high mRNA levels in a previous STARR-seq experiment.

#### A.1.2 Design of synthetic (non ML) 3’ UTRs for training data curation

To design a set of synthetic UTRs to use as training data, we collected motifs suspected to affect mRNA stability. We first searched for motifs based on the results of our high-throughput stability assay of our initial genomic 3’ UTRs and control 3’ UTRs. We selected the 130 most stable 3’ UTRs and compared them to 200 background sequences selected from our dataset, using MEME to identify motifs that were more common in our stable 3’ UTRs [46]. We additionally extracted known activating RNA binding protein (RBP) motifs from [47] and oRNAment [48]. As a control, we included de-stabilizing motifs, including de-stabilizing RBPs and AU-rich elements (ARES). Motifs were clustered and merged to reduce redundancy using gimme motifs [40].

We next collected a set of RNA secondary structural elements from internal and external data sources. We first identified de novo structural motifs in stable genomic sequences using BEAM, a secondary structure motif discovery tool [49]. We additionally include structural motifs from and RNAVlab [50].

Finally, all structural and canonical motifs were randomly sampled and inserted into 210 stable 3’ UTRs chosen from our existing stability assays. We inserted single motifs into backbones (1 to 6 times), pairs of motifs into backbones, and any number of 3 to 6 randomly chosen motifs into backbones.

#### A.1.3 Additional genomic and synthetic (non ML) 3’ UTRs

Additional genomic sequences were selected from a variety of sources, including high stability human and mouse 3’ UTRs [47, 51], flaviviral 3’ UTRs, Thermus thermophilus RNAs, and human structural RNAs. These genomic sequences were tested in later iterations and were not included in our initial 3’ UTR dataset.

**Figure A4:**
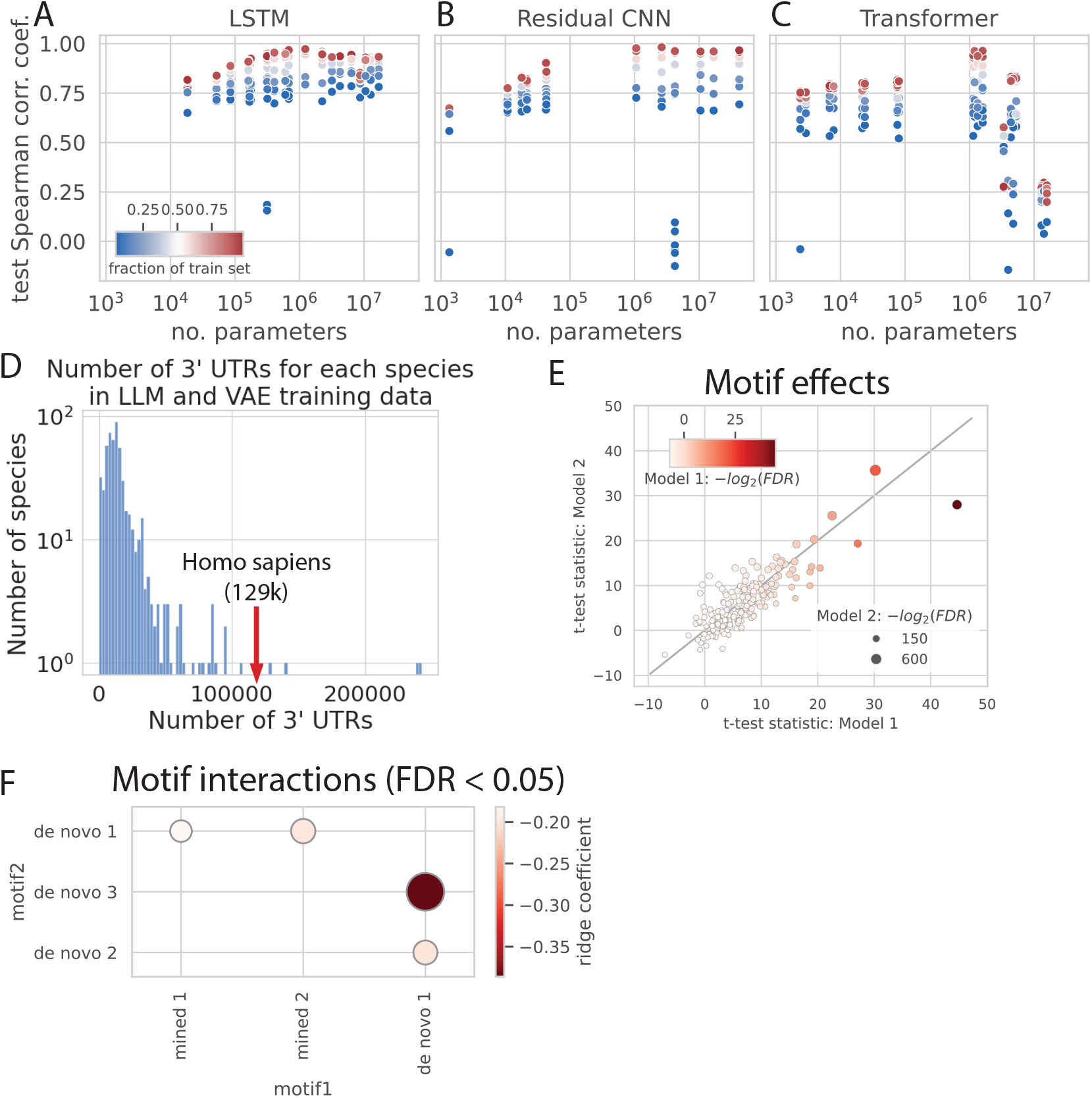
Comparison between number of model parameters and Spearman rank correlation coefficient of a held out clustered test set for (a) LSTM (b) Residual CNN (c) transformer architectures (d) Histogram of numbers of unique 3’ UTRs in training dataset used to train LLMs and VAEs for each species. 129,473 unique 3’ UTRs from homo sapiens are labeled. (e) Significance and effect size of 212 motifs present in 3’ UTRs and their effect on predicted mRNA stability. Plot compares predicted significance for each motif across two model architectures. Test for significance was performed for each motif by placing motifs in randomer sequences, and comparing predictions between randomers with motifs and a dinucleotide shuffled set. (f) Significance and effect size of motif pairs present in 3’ UTRs and their effect on predicted mRNA stability. 212 motifs were inserted at random into a neutral randomer sequence. Ridge regression was performed on sequences, using the presence of all motif pairs as features and predicted stability as a response variable. Four significant interactions were identified (FDR < 0.05), which includes two motifs that were mined from public datasets and three motifs that were identified from models using TF-MoDISco [34].

In iteration three we generated additional synthetic, non-ML synthetic sequences as shown in Figure 2 c. One set of 3’ UTRs was generated by removing and adding miRNA binding sites into a range of backbones. Another set was generated by concatenating tiles together from 3’ UTRs that were measured to be stable in previous iterations of design. Tile sizes were 62bp, 83bp, and 126bp. For a subset of these designs, we filtered them using a supervised model. If the model was used for filtering, the sequences were categorized as ML designs.

### A.2 MPRA stability assay

#### A.2.1 Library construction

Libraries were ordered as ssDNA oligo pools from Twist Bioscience, with left and right flanking adapters for cloning. Upon arrival, oligo pools were amplified with primers binding to the adapters and adding 15-30 base pairs of homology to CDS and 3’ constant region of the backbone plasmid, respectively, for cloning via Gibson assembly.

Backbones were constructed in an ampicillin-resistant pDonor vector to include the T7 promoter, hHBB 5’ UTR, CDS of choice, AgeI and PmeI recognition sites, and 3’ constant region. hHBB 5’ UTR was selected due to hHBB’s efficient expression in mammalian mRNAs [52, 20, 21]. Backbone plasmids were digested with AgeI and PmeI for insertion of amplified oligo pools.

Gibson Assembly was performed by mixing 50 ng of linear backbone and 2:1 molar excess of amplified oligo pools with 2x NEBuilder HiFi DNA Assembly Master Mix. Reactions were cleaned up with Monarch PCR & DNA Cleanup Kit and electroporated into NEB 10-beta competent E. coli.

#### A.2.2 mRNA synthesis

*In vitro* transcription (IVT) templates were prepared by digesting 25 µg plasmid DNA with MfeI-HF at 37C for 2 hours. Reactions were purified by phenol-chloroform extraction and tested by Nanodrop and E-Gel for concentration, purity, and complete linearity. IVT was performed using standard nucleotides or N1-methylpseudouridine as noted. A ∼ 150 base poly A tail was added via A-Plus Poly(A) Polymerase Tailing Kit and a 5’ cap was added using CleanCap Reagent AG.

#### A.2.3 MPRA experiment

IVTed mRNA libraries were transfected into HEK293T cells in triplicate using either lipofectamine or electroporation methods. Additional experiments used HeLa cells, hSkMC cells, and *in vivo* mouse muscle. Following transfection, RNA was extracted from cells at 1, 3, 24, 48, and/or 72 hour time points. Total cellular RNA and MPRA mRNA were extracted together from cells following standard RNA isolation protocols with TRIzol and Qiagen RNAeasy Mini Kit.

### A.2.4 Library preparation and sequencing

Library prep consisted of reverse transcription, qPCR, full-length enrichment PCR, and an indexing PCR. RNA input for reverse transcription was based on estimated amounts of MPRA mRNA in the samples at each time-point, rather than cellular RNA measurements. RT was carried out with SuperScript IV Reverse Transcriptase and primer binding to 3’ constant region and adding a 16bp unique molecular identifier (UMI). cDNA was cleaned up with Monarch PCR & DNA Cleanup Kit. qPCR was performed to determine the number of unique full-length mRNA molecules per sample. For consistency in PCR cycles and sequencing depth, the same number of cDNA molecules were carried forward to PCR for each sample. The first round of PCR (10-20 cycles) used primers set at the 5’ and 3’ ends of the molecule to enrich for full-length transcripts, as described by Leppek et al [20]. After a 1X Kapa bead clean-up, an additional 5 cycles of PCR were performed using primers that add Illumina P5 regions, i7 indices, and Illumina P7 regions. Final libraries were run on 4% E-Gel and extracted using Monarch DNA Gel Extraction Kit Protocol. Libraries were quantified by Qubit 1X dsDNA BR Assay and pooled in equal molar amounts for sequencing on Illumina MiSeq or NextSeq.

#### A.2.5 Data processing

A customizable nextflow pipeline was designed to process Illumina sequencing read data into 3’ UTR counts and half-lives across time point samples. Reads were aligned to library reference sequences using Burrows-Wheeler Aligner bwa-mem [53] to determine their 3’ UTR of origin. For each sample, read UMIs were grouped and deduplicated using umi_tools [54], and UMIs were further cleaned using a custom python script to remove errant UMIs (mononucleotide UMIs and UMIs matching adapter sequences) and UMIs shared between 3’ UTRs. Custom quality control R scripts further examined reads per UMI and total UMIs per sample and per 3’ UTR.

#### A.2.6 Calculation and normalization of half-lives

For each assay we infer half-lives of different mRNA species from experimental next generation sequencing (NGS) count data sampled over multiple time points. We construct a statistical model of degradation that assumes that mRNA decays exponentially over time. Half-lives can be modeled as a truncated normal distribution, with mean priors estimated empirically from normalized UMI counts for each UTR and standard deviation prior as a half normal distribution. Observed UMI counts are modeled as a multinomial distribution, which is parameterized by the log degradation rate of each candidate. Inputs to the model are a count matrix, the time points and replicate relationships, and any known half-lives for included benchmarks. The model is specified as follows.

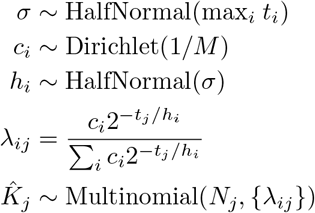

In this model, 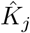 is the observed UMI count vector for sample *j. λ*_*ij*_ is the expected fraction of UMIs that come from candidate *i* when randomly sampling from sample *j*. (It has been adjusted accordingly for the expected degradation over time.) Note that Σ_*i*_ *λ*_*ij*_ = 1, so they are used as multinomial parameters when drawing observed UMI counts. Note how we normalize the underlying relative amounts in *λ*_*ij*_. This is because the starting concentrations and half-lives are independent, and rapid degradation of one candidate should increase the relative concentration of another as time goes on. *M* is the complexity of the library. *N*_*j*_ is the total number of UMIs in sample *j. t*_*j*_ is the time point for sample *j* (measured in hours, for example). *c*_*i*_ is the underlying (initial/background) concentration of candidate *i. h*_*i*_ is the half-life of candidate *i*.

We also include a likelihood term for the “observed” half-lives to help enforce agreement between the inferred *h*_*k*_ and the prior mean/uncertainty of the benchmarks 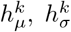 for benchmark *k*:

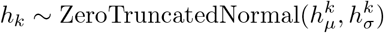

The inference is implemented with the NumPyro probabilistic programming framework [55]. We used Markov chain Monte Carlo (MCMC) to sample from the posterior distribution. MCMC was run for 1,000 samples, with three chains and 3,000 warmup steps.

To normalize half-lives across assays for curating a single training dataset, half-lives were z-score transformed using the mean and standard deviation of half-lives from 3’ UTRs that were included as shared carryovers in all assays. Z-score normalized half-lives were then re-transformed back to a single reference assay to rescale values within the range of half-life in hours. While ML models were trained on z-score normalized half-lives (shown in Figures 4 b and 2 d), measured half-life values (depicted in Figure 2 e, Supplementary Figure A2 cd) depict rescaled half-lives.

### A.3 *in vivo* validation

We selected three high stability 3’ UTRs to compare to the hHBB control 3’ UTR *in vivo*. 3’ UTRs were placed after a hHBB 5’ UTR and a luciferase coding sequence. mRNA was synthesized via IVT and then formulated with MC3 into lipid nanoparticles (LNPs) by GenScript. These mRNA-LNPs were then sent to Biomere for *in vivo* animal studies. mRNA-LNPs were systemically delivered to five C57BL/6J mice each via lateral tail vein injection at a dose of 0.5 mg/kg. Protein activity was measured daily via *in vivo* imaging system (IVIS) imaging – luciferin was delivered systemically, and luciferase activity was quantified by bioluminescence quantification in total flux (photons/second) of the ventral and dorsal animal images.

We observed outlier animals and outlier samples in the returned data, which we removed. Four animals (two in the hHBB group and two in the TC9 group) were removed due to exhibiting 5-10x reduced fluorescence throughout the experiment compared to other animals in their respective groups. We also observed “drop out” luminescence measurements at later time points for specific animals (TC9 animals 2 and 3 on Day 5 and TC11 animal 3 on Day 2), though preceding and following days showed comparable luminescence to other animals. We hypothesized that those dropout measurements were due to luciferin injection errors and removed them from further analyses.

Total flux measurements were used to estimate relative protein expression, total protein production (area under the curve), and mRNA half-life. Due to variance in initial protein production and the presence of outlier animals, we normalized relative protein expression at each time point to the first time point (hour 6) for each animal, and we then compared each animal to the average relative protein expression of the hHBB control animals (Fold change vs hHBB, normalized to 6hr). We calculated total protein production per animal using an area under the curve calculation of the total flux measurements, assuming log-linear decay between time points. We also calculated the relative total protein production per animal using the normalized relative protein expression curves (Fold change vs hHBB, normalized to 6hr), again assuming log-linear change between time points. Finally, we calculated mRNA half-life per animal. Given the short half-life of luciferase protein, we determined that the protein abundance measurements could be used to approximate the current mRNA levels. We selected measurements from day 1 to day 4, before any animals reached our lower limit of detection (<5×10e5), and fit a linear model to the log2-transformed protein measurements. We took the negative inverse of the linear model slope to approximate the mRNA half-life, i.e. the time it takes for mRNA to reduce by one unit on the log2-scale.

### A.4 Generation of train, validation, and test data splits for evaluating supervised models

To generate a test set that was distinct from the training distribution, sequences were first clustered using mmseqs [56] using easy-cluster and the following parameters: “-c 0.1 -s 7.5”. All clusters that contained at least one very stable UTR (defined as z-score transformed half-life > 5) were reserved for the test set. Holding out all entire clusters that contain the most stable UTRs allow us to evaluate whether models can generalize to predict stability of UTRs that exceed stability in the training set.

To sample training and validation splits for evaluation of architecture performance in Figure 4 a, we randomly sample 5% to 98% of clusters (including all sequences in a selected cluster), and randomly select 15% of the selected sequences for a validation set, regardless of cluster assignment.

### A.5 Curation of 3’ UTRs from 550 eukaryotes for training unsupervised models

To train unsupervised 3’ UTR variational autoencoders and language models we assembled a collection of 3’ UTRs from UTRdb 2.0 [18], which contains 5’ and 3’ UTRs from eukaryotic mRNAs. We pulled all 3’ UTRs from 550 species that were shorter than 5,000 nucleotides. Duplicate sequences, sequences with invalid characters, and sequences shorter than 20nt or longer than 5000 nt were discarded. After filtering, 9.5 million unique sequences remained for training.

We additionally filtered 125 of the 550 species that were specifically mammalian to curate a mammalian training dataset. This dataset consisted of 1.9 million 3’ UTRs.

UTR training data splits and species metadata used for unsupervised models are publicly available at Google Drive.

### A.6 Supervised model architecture

Figure 4 a demonstrates three supervised architectures that were compared for evaluating predictive accuracy of 3’ UTR stability. Input to all supervised architectures is the one-hot encoded representation of the 3’ UTR. The loss function is mean squared error (MSE), as the task of predicting mRNA stability from 3’ UTR sequences is treated as a regression problem. Models were trained for a maximum number of 1,000 epochs with early stopping based on the validation loss to prevent overfitting. Models were trained via Pytorch Lightning [57] using the Adam optimizer with an initial learning rate of 1e-3, 1e-4 weight decay, and default betas (0.9, 0.999). Learning rates were reduced upon plateauing based on validation loss. All models were trained on a single NVIDIA A100 GPU on a2-highgpu-8g machines.

#### A.6.1 LSTM architecture

We implemented a Long Short-Term Memory (LSTM) architecture to predict mRNA stability from 3’ UTR sequences. This architecture first processes sequences through a single convolutional layer with kernel width of 10 with a stride of 1 and no dilation. Features from the convolutional layer are passed through ReLU activation and max pooling along the sequence dimension with kernel width of 5 and stride of 3. Output from max pooling is passed to a bidirectional LSTM, then passed through a single linear layer. We use a dropout layer with a dropout rate of 0.5 before the LSTM to prevent overfitting. In Figure 4 a we vary the following hyperparameters to test the effect of model size on test set accuracy: the number convolutional filters (ranging from 32 to 1024) and number of features in the LSTM’s hidden state (ranging from 32 to 1024). The number parameters for all models shown in Figure 4 a range from 18,273 to 16,837,633, based on the chosen hyper parameters.

The selected LSTM that was used to generate stability predictions in Figures 4 b-e and Supplementary Figures A4 e-f have 256 convolutional filters and 32 features in the LSTM’s hidden state, with a total of 84,801 parameters.

#### A.6.2 Residual Convolutional architecture

We additionally implemented a residual convolutional architecture to predict mRNA stability from 3’ UTR sequences. This architecture consists of multiple convolutional blocks. Each convolutional block consists of a convolutional layer and a ReLU activation layer, where a residual connection is added between the output of the activation layer and the original inputs to the block. Outputs of the last convolutional block are fed through a max pooling layer and a linear layer. In Figure 4 a we vary the following hyperparameters to test the effect of model size on test set accuracy: the number of convolutional filters (ranging from 32 to 1024), and the number of convolutional blocks (ranging from 1 to 6). The number parameters for models shown in Figure 4 a ranged from 1,345 to 41,991,169, based on the chosen hyperparameters.

#### A.6.3 Transformer architecture

Similar to work in [58] we built on a transformer architecture which first processes sequences through a single convolutional layer with kernel size of 10. Positional embeddings are added to outputs of the convolutional layer and are passed to a transformer encoder layer, with ReLU activation, 8 attention heads, and 0.1 dropout. Outputs from the final transformer block are passed through two linear layers (as in [59]). In Figure 4 a we vary the following hyperparameters to test the effect of model size on test set accuracy: the number of convolutional filters (ranging from 32 to 1024), the number of transformer blocks (ranging from 1 to 3), and dimensions of the feed forward layer in each transformer block (ranging from 16 to 512). The number parameters for models shown in Figure 4 a range from 2,385 to 15,801,857, based on the chosen hyper parameters.

### A.7 Variational Autoencoder architecture and training

Model architecture. We trained two variational autoencoders (VAEs) on 3’ UTRs from 550 eukaryotic species and 125 mammalian species, respectively. Both eukaryotic and mammalian models used a transformer encoder/decoder architecture with a parameter size of 5.7M. Both the encoder and decoder were autoregressive transformers based on the original Transformer architecture [59]. While inputs to the encoder include positional embeddings and 3’ UTR sequences that were converted representing the four nucleotides, inputs to the decoder include a latent vector from the encoder of dimension 256, in addition to positional embeddings. Both the encoder and decoder have three transformer layers. Each layer consists of self attention with four attention heads and a feed forward network. Feed forward dimensions for the encoder and decoder were 256 and 128, respectively. The latent dimension of encoder outputs and corresponding decoder inputs is 256.

#### Model loss

The loss is computed from the reconstruction error of each 3’ UTR and the Kull-back–Leibler (KL) divergence between the learned latent distribution and a prior distribution, which we define as a Gaussian distribution. The total loss for the VAE is formulated as in [60]:

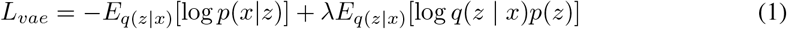

Where x is the input UTR sequence, z is the latent variable, *p*(*x* | *z*) is the probability of reconstructing *x* given the latent representation *z, p*(*z*) is a unit Gaussian prior, and *q*(*z* | *x*) is the approximate posterior distribution. While the first position of this equation defines the reconstruction loss, the second position defines the KL divergence loss. We set *λ* to 0.1 during training.

#### Training details

During training, we used early stopping based on validation loss to prevent overfitting. Both encoders and decoders used a linear warmup schedule for learning rates. The decoder learning rate was set to 0 until encoder warmup steps were completed.

#### Generating UTR samples

Latent variables were first generated by sampling from a Gaussian prior. Due to the autoregressive design of the decoder, a start token was initially passed through the decoder, along with the sampled prior as memory. The next token was subsequently generated by sampling the output of the decoder for each position and generating the selected token using Gumbel-Softmax [61]. This procedure was run in an autoregressive fashion until a stop token was generated.

### A.8 3’ UTR masked language architecture

We trained a 44M parameter masked language model on mammalian 3’ UTRs. This dataset was the same dataset used to train the mammalian VAE, and is described in A.5. We trained Bidirectional Encoder Representations from Transformers (BERT) [62] with a maximum context length of 1000 on single nucleotide tokens from UTRs. The BERT model included six hidden layers, eight attention heads, with a maximum sequence length of 1000. Models were trained using a masked language model objective, masking UTRs at 15% during training. Models were trained using the BERT architecture available in the Hugging Face interface[63].

BERT models shown in Figure 4 a were fine tuned on 3’ UTRs and corresponding stability measurements using a learning rate of 5e-5, mean squared error loss, and a linear learning rate warmup of 100 steps.

#### A.8.1 Statement of model access and responsibility

Our 3’ UTR masked language model is available for querying masked sequences and collecting model embeddings from 3’ UTR sequences at https://models.ginkgobioworks.ai/models. We believe that we are responsibly releasing this model because:

1. The published model was only trained on publicly available, genomic sequences (no synthetic sequences).
2. The published model is only trained on DNA sequence (no functional information). Therefore, we do not anticipate that users can directly leverage this model to design or modify existing sequences in a way that directly optimizes the sequence’s functionality.
3. The provided API does not allow access to the model weights or gradients.

## A.9 Simulating individual motif effects and motif interactions using a predictive model

Figure 4 e demonstrates the significance and effect size of each motif towards increasing the predicted stability of a 3’ UTR when compared to a set of dinucleotide shuffled sequences. The following procedure was performed for 212 motifs: 1000 samples were sampled from a motif and were embedded at a randomly chosen position of a randomly generated DNA sequence of length180. Each generated sequence was scored using two ML models and compared to 1000 sequences that were dinucleotide shuffled from the generated sequences. We use two models, as displayed in Figure 4 e, to ensure that significance and effect size of motifs correlates across models. A t-test was performed to determine whether predictions for randomers containing a motif were significantly greater than dinucleotide shuffled sequences that broke the motif. Corresponding p-values were adjusted using false discovery rate across all motifs.

Figure 4 f demonstrates the significance and effect size of pairs of motifs that were predicted with our model. To identify predicted motif interactions, we first generated a training dataset of 10,000 sequences that randomly inserted 1 to 5 sampled motifs into a neutral randomer sequence.These 10,000 sequences were split into a training and validation set, and ridge regression was performed on extracted sequence features that included the presence of each motif pair and a response variable of predicted stability of each sequence. P-values were calculated for each ridge regression coefficient and were adjusted using false discovery rate across all motifs pairs. Motif pairs with FDR < 0.05 were considered significant.

### A.10 ML Design Methods

#### A.10.1 Genetic algorithm

We implemented a genetic algorithm inspired by [14]. An initial set of stable 3’ UTRs were selected as seed sequences to initially pass to our genetic algorithm. We ran a genetic algorithm for 20 iterations. In each iteration, seed sequences were randomly recombined with an average recombination size of 50 nucleotides. Recombined sequences subsequently underwent 10 rounds of mutations, where five mutations were added at random to each sequence and were then sampled weighted by predicted stability. At the end of each iteration, original seed sequences and the recombined and mutated sequences were sampled from to generate the next iteration of seed sequences. Original seed sequences were sampled from with 5 times the probability of the mutated sequences. The number of sequences selected during sampling was determined by Vendi score diversity (calculated from kmers) [22], where more diverse sets were allocated larger sample sizes.

During each mutational round, mutations were chosen either (1) at random positions, (2) based on the predictions from a supervised stability model, or (3) using likelihoods from a 3’ UTR language model. The effect of the method for mutation selection on measured stability is shown in Supplementary Figure A2 c. To select mutations based on a supervised stability model, we ran *in silico* mutagenesis on each sequence and selected all mutations with a mutation effect >0.1 when compared to the predicted stability of the original sequence. To mutate based on likelihoods using a 3’ UTR language model, sequences were randomly masked and fed through the language model (See A.8). For each masked position, the nucleotide with the highest predicted likelihood was selected for mutation.

#### A.10.2 Simulated annealing

Simulated annealing was run with the same format and parameters as our genetic algorithm. However, recombination and mutations were selected based on an annealing temperature that decreased exponentially based on the number of iterations.

#### A.10.3 Seqprop

We employed an implementation of activation maximization, proposed in SeqProp [64]. Our implementation of SeqProp utilizes backpropagation to iteratively optimize parameters of a matrix that is sampled from to generate 3’ UTRs. Generated 3’ UTRs are passed through a pre-trained predictive model that is frozen during training. During training, loss is computed from the negative predicted stability of the generated UTR, and is regularized by the entropy of the samples to increase diversity of generated sequences. Hyperparameters include a learning rate of 0.001, trained for a maximum of 1000 epochs. Early stopping is implemented to stop training when predicted stability converges to a local minima.

#### A.10.4 Conditioning by Adaptive Sampling (CbAS)

We implement CbAS [17] by first training a variational encoder on genomic 3’ UTRs to learn the distribution of natural sequences (See A.7). For 50 iterations, we sample from a VAE and select the top 30% UTRs with the highest predicted stability (predicted from a supervised stability model), and regularize selection of UTRs based on the KL divergence between the original and tuned VAE. A VAE is re-trained on selected UTRs, and iteratively re-sampled to generate UTRs that further increase predicted stability. Regularization during sequence selection inhibits the distribution learned by the tuned VAE to diverge significantly from the original VAE.

#### A.10.5 Design by Adaptive Sampling (DbAS)

DbAS [16] was run with the same models and parameters as CbAS, except the regularizing VAE was removed, allowing for samples produced from the tuned VAE to exit the sequence distribution learned by the original VAE.

#### A.10.6 Synthetic (ML selected)

Synthetic sequences were generated by placing structural and canonical motif samples in various 3’ UTR backbones as discussed in A.1.2. Synthetic sequences were rank ordered by predicted stability, and sequences with the highest predicted stability were printed in an oligonucleotide pool and tested in a stability MPRA.

#### A.10.7 *insilico* Mutagenesis (selected with predictive model)

We ran *in silico* mutagenesis on a set of stable 3’ UTRs. For each mutated position and each sequence, we calculated the mutation effect based on the difference between the predicted stability of the mutant sequence and the predicted stability of the unmutated sequence. All mutants that had a mutation effect > 0.1 were included in the final mutated sequence. Levenshtein distance was computed between unmutated and mutated sequences, whose distances ranged from 1 to 58 nucleotides.

#### A.10.8 *in silico* mutagenesis (selected with masked language model)

A set of stable 3’ UTRs were masked at various masking probabilities (ranging from 6% to 60% masking) and fed through a BERT language model trained on mammalian 3’ UTRs (See A.8). For each masked position, the nucleotide with the highest predicted likelihood was selected for mutation with respect to the original 3’ UTR. Levenshtein distance was computed between unmutated and mutated sequences, whose distances ranged from 1 nt to 39 nt.

### A.11 Testing the effects of active learning on model performance

We evaluated three active learning methods: core-sets [25], margin-sampling [26], and cluster-margin [27]. Each method selects potentially informative sequences from a large candidate pool, aiming to enhance model performance by training on the selected sequences. Core-sets prioritize selecting a diverse set of sequences, while margin sampling focuses on sequences that our models are most uncertain about. Cluster-Margin aims to choose diverse sequences that the model is also uncertain about. Our candidate pool consisted of approximately 15 million synthetic 3’ UTRs generated by inserting motifs into 3’ UTR backbones as described in A.1.2. We used an ensemble to predict the stability of each 3’ UTR and also gathered the variance in predictions across constituent models as a measure of model uncertainty. Out of the 15 million initial candidates, we filtered out sequences with low predicted stability to obtain 1 million final candidates for active learning-based selection. Then, each method was used to select 2,500 UTRs. Core-sets and Cluster-Margin need pairwise distance metrics to perform diversity-based selection. To obtain these distances between sequences, we performed PCA reduction (dimensions = 256) on model embeddings for candidate sequences to obtain a vector representation for each sequence. Then, a standard euclidean distance metric was computed between sequences. Finally, as a baseline, we randomly sampled UTRs from the set of 1 million sequences, weighted by predicted stability.

Experimental measurements for active learning sequences were collected and filtered for high quality measurements, in which we retained between 1400 to 1700 UTRs for each method. We retrained four models on the original training set used when performing active learning selection (65,200 sequences), as well as the additional 3’ UTRs collected from each active learning method. Final performance of each active learning method was evaluated as the Spearman correlation coefficient on the test set of the original model used during design of the active learning UTRs.

### A.12 Selection of three ML designed 3’ UTRs for *in vivo* validation

Three candidate 3’ UTRs were selected for *in vivo* validation based on quality filters and measured half-lives across assays three of distinct assays. Quality filtered include low standard error across replicates and sufficient reads per UMI at 24 and 48 hr samples. Sequences were further filtered based on their half-life improvement relative to the hHBB 3’ UTR benchmark. Additionally, we only considered UTRs whose improvement over hHBB 3’ UTR increased between 24 and 48 hours. These filters resulted in 19 distinct sequences, which were further filtered based on sequence diversity and the absence of muscle miRNA and monocyte-derived dendritic cell binding sites.

